# MMR Deficiency Defines Distinct Molecular Subtype of Breast Cancer with Unique Proteomic Networks and Variable Clinical Significance

**DOI:** 10.1101/2022.04.14.488341

**Authors:** Sean M. Hacking, Charissa Chou, Yigit Baykara, Yihong Wang, Alper Uzun, Ece D. Gamsiz Uzun

## Abstract

Mismatch repair (MMR) alterations are important prognostic and predictive biomarkers in a variety of cancer subtypes including colorectal and endometrial. However, in breast cancer (BC), the distinction and clinical significance of MMR is largely unknown. This may be due in part to the fact that genetic alterations in MMR genes are rare, and only seen to occur in around 3% of BCs. In the present study we analyzed TCGA data using a multi-sample protein-protein interactions (PPI) analysis tool, Proteinarium, and showed a distinct separation in the MMR deficient and intact specific networks. MMR deficient tumor specific networks have a highly connected cluster of histone genes, identified by unique PPI. We also found that distribution of MMR deficient breast cancer is more prevalent in HER2-enriched and triple-negative (TN) BC subtypes compared to luminal BCs. Poorer survival was seen in patients with HER2-enriched BCs with MMR deficiency, whereas an improved survival was seen in TNBCs with MMR deficiency. We recommend defining MMR deficient breast cancer by next generation sequencing (NGS) when any somatic mutation is detected in one of the 7 MMR genes found in our study. Our recommendations include labeling patients with variants of undetermined significance (VUS) as MMR deficient supported by findings from distinct clusters of patients based on our network analysis. MMR may have a role in guiding the use of immunotherapy for both TN as well as HER2-enriched BC.

## Introduction

Worldwide, a woman is diagnosed with breast cancer (BC) every 9 seconds^1^. Mismatch repair (MMR) is important in daily clinical practice for the cancer management in a variety of cancers including colon and endometrium as screenings are used to identify for the potential of Lynch syndrome^2^. Today, the role of MMR in breast cancer is obscure and not well understood. An early report^3^ in 16 BC patients demonstrated no incidence of MMR in five microsatellite loci (D2S123, D3S1611, D17S807, D17S796 and Xq11-12), initially suggesting MMR to be uncommon in human early-onset breast cancer and not appear to be related to prognosis. In BC most clinical emphasis is placed on a patient’s individual hormone receptor (HR) status: estrogen receptor (ER), progesterone receptor (PR), and human epidermal growth factor receptor 2 (HER2); as well as tumor proliferative index measure by Ki-67. Roughly, this allows tumors to be subdivided into the following classical molecular subtypes: Luminal A (ER+/−, PR+/−, HER2-), Luminal B (ER+/−, PR+/−, HER2+, or Ki67>14-20%), HER-2 positive (ER-, PR-, HER2+), and triple negative (TN)/basal (ER-, PR-, HER-2-)^4^. A normal-like subtype has also been described, although its significance is still unclear, they are characterized by normal breast tissue profiling^5^.

MMR encompasses the post-replication preservation of DNA homeostasis and genomic stability^6^. The MMR system functions not only to correct spontaneous base–base mispairing, but also small insertions–deletions (indels) that occur following DNA replication; subsequent failure results in an increased mutation rate^6^. The MMR system contains a group of genes which encode proteins interacting as heterodimers. In humans there are 8 genes which make up the MMR machinery and encode for its protein components. These include the homologs of *E. coli* MutS genes: MSH2, MSH3, MSH5, and MSH6, and the MutL genes: MLH1, PMS1 (MLH2), MLH3, and PMS2 (MLH4)^6–8^. Mutations in MMR can lead to a loss of translation and subsequently genomic instability, however MSH5 has not been found to be associated with MMR in humans^6^. MSH2, is located on chromosome 2p21 and is considered the principal MSH protein which creates distinct heterodimers with MSH6 and MSH3, referred to as MutSα and MutSβ, respectively^9^. MLH1 encodes a protein that dimerizes with PMS2 to form MutLα, and MLH3 which forms MutL*β*^9^.

Most recently, genome-wide association studies (GWAS) have grown in popularity as it offers an approach to investigate complex diseases such as BC^10^. However, GWAS has fallen short in identifying ‘missing heritability’, our fundamental understanding of the complex architecture of the genome due to additive genetic effects is not fully represented by GWAS^11^. Analyzing the complex architecture could demonstrate subgroups of patients which share variants in genes in specific networks with shared phenotype. To combat this problem, we recently developed Proteinarium, a multi-sample protein-protein interaction (PPI) tool with the ability to identify clusters of cancer patients with shared gene networks^11^. Proteinarium functions by converting user defined seed genes into corresponding protein symbols, which is then mapped onto the STRING database’s interactome^12^. A PPI network is built for each sample using Dijkstra’s algorithm^13^. Pairwise similarity scores are calculated to compare the networks and cluster the samples where they are presented as a dendrogram. A layered graph of PPI networks for the samples in any cluster can be visualized and analyzed.

The principle aims of this study are to determine whether MMR deficient BC is associated with a distinct molecular subtype, identify unique PPI networks, and finally compare the clinical significance as well as prognostic value of MMR across molecular BC subtypes. This could allow for a better understanding, demonstrate clinical significance, and facilitate a standardized definition of MMR deficiency in BC.

## Material and Methods

### Study Design

This study included cases from the Breast Invasive Carcinoma (TCGA, PanCancer Atlas)^14–23^. The RNA-seq data was obtained from the cBioPortal (https://bit.ly/3g4lTM4). Tumor data was obtained from patients that had not received prior treatment for their disease (ablation, chemotherapy, or radiation therapy). The initial cohort included 1084 patients. Cases without any genomic data in the following genes were excluded: MSH2, MSH3, MSH6, MLH1, PMS1, MLH3, and PMS2. Final cohort included 996 patients.

### Proteinarium

Proteinarium, a novel multi-sample tool used to identify clusters of patients with shared PPI networks, was used to compare the MMR deficient and MMR intact patients. MMR deficiency was defined as having a mutation in one or more of the eight MMR machinery genes. The seed genes – top 100 upregulated genes – for each patient were selected using z-score data from the RNA-seq data. Proteinarium uses the experimentally validated PPI information from the STRING Database to build network graphs based on the seed genes for each patient. The similarity between each pair of network graphs is calculated using the Jaccard distance, which is recorded in a similarity matrix used as an input for clustering graphs. Unweighted Pair Group Method with Arithmetic Mean (UPGMA) is used to cluster patients, with their network similarities as an output. Significance of a cluster is calculated using Fisher exact tests, based on the abundance of cases and controls. The minimum path length was set at two, and thus included only the pathways of seed proteins connected directly to each other and/or via a single intermediary protein (“imputed genes”). Because the number of MMR intact and MMR deficient patients was widely unequal (89 MMR deficient versus 907 MMR intact patients), we performed ten iterations of Proteinarium runs comparing 89 MMR deficient patients versus 90-91 randomly selected MMR intact patients. The output of each Proteinarium run was analyzed for the presence of significant clusters (p-value < 0.05) dominated by MMR intact patients. The MMR intact patients in the cluster with the largest number of MMR intact patients and no more than 2 MMR deficient patients (if there were no clusters with 0 or 1 MMR deficient patients) were compiled in a list of “selected” MMR intact patients. These MMR intact-dominated clusters had patients with significant PPI network similarity. After running ten iterations of the randomly selected MMR intact vs. MMR deficient patients in Proteinarium, the list of selected MMR intact patients contained 67 patients. Finally, Proteinarium was used to compare the 67 selected MMR intact patients and 89 MMR deficient patients. This yielded a dendrogram and list of significant clusters that was used for further analysis.

### Separation Test

Separation testing was used to determine the genetic similarity between diseases or phenotypes of diseases by comparing their PPI networks in the interactome. We performed a separation test to determine the distance between the PPI networks of the MMR deficient and MMR intact dominated clusters using a Python script adapted from Menche et.al.^24^. The interactome is composed of 141,296 experimentally determined physical interactions between 13,460 proteins. s_AB_ is a measure of the distance between two diseases in the interactome, with a positive value indicating a topological separation of the disease modules and a negative value indicating that the diseases overlap in the interactome.

### Statistical Analysis

Means and frequencies were calculated for all variables using Microsoft Excel (*Version 16.58*). Comparative analysis and Kaplan-Meier survival analysis were performed using variable statistical methods provided in the cBioPortal user interface. Kaplan Meier survival plots were made to demonstrate survival curves in relation to time in months by the log rank methods. Comparative analysis was performed using the non-paired t-test, Fisher’s Exact test and chisquare contingency test to compare the MMR deficient dominant cohort (cluster 71) with control MMR patients. These statistical analyses were performed using Prism Graphpad version 8.4.2 and a p-value < 0.05 was considered statistically significant.

## Results

### Mutational Portraits of MMR Genes Signatures in Human Breast Cancer

Out of the 1084 patients in the Breast Invasive Carcinoma (TCGA, PanCancer Atlas), 996 patients were included based on availability of sequencing data and 89 were found to have mutations, structural variants or copy-number alterations in any of the 7 MMR genes: MSH2, MSH3, MSH6, MLH1, PMS1, MLH3 and PMS2. No mutations were seen in MSH5. Mutations included both driver as well as variants of undetermined significance (VUS). MSH2 had a somatic frequency of 1.0% with 4 driver mutations: 1 truncating, 1 splice, and 2 SV/fusion; 6 VUS, all of which were missense. MSH3 had a somatic mutation frequency of 0.7% with 3 driver mutation, all truncating, and 5 VUS, all of which were missense. MSH6 had a somatic mutation frequency of 0.9%: 2 truncating and 1 SV/fusion; 8 missense variants which were VUS. MLH1 had a somatic mutation frequency of 0.9% with 5 driver mutations: 1 missense, 2 truncating, 1 spice and 1 SV/fusion; and 5 VUS, all of which were missense. PMS1 had a somatic mutation frequency of 1.2% with 3 driver mutations: 2 truncating and 1SV/fusion; 9 VUS were seen, all missense. MLH3 had a somatic mutation frequency of 1.1% and contained 12 VUS, 11 of which were missense and 1 inframe. PMS2 had a somatic mutation frequency of 0.8% and contained 2 driver mutation, all truncating: and 6 VUS, all missense. 3D protein structural modeling for mutations in 7 MMR genes can be seen in Supplemental Figure 1.

### MMR Deficient and Intact Patients have Distinct PPI Networks

The Proteinarium dendrogram output revealed significant clusters (p-value <0.05) dominated by either MMR deficient or MMR intact patients. Two clusters were chosen for further investigation based on degree of domination by either MMR deficient or MMR intact patients and number of patients in the cluster (Figure 1). The MMR deficient cluster contained 52 MMR deficient patients and 0 MMR intact patients while the MMR intact cluster contained 51 MMR intact patients and 6 MMR deficient patients. The consensus networks for significant clusters dominated by MMR deficient and MMR intact patients include genes unique to each MMR status group. DMP1 was found to be the hub gene present only in MMR intact patients. DMP1 is an extracellular matrix protein critical for the proper mineralization of bone and dentin. Of note in the MMR deficient consensus network was the presence of a subnetwork of five histone genes, four of which are from the histone H2B family and were unique to MMR deficient patients. The one gene encoding a member of the histone H2A family was imputed. GNB1, which encodes the beta subunit of G proteins, and CDK1, encoding the catalytic subunit of the M-phase promoting factor necessary for G1/S and G2/M phase transitions in the cell cycle, are both highly connected genes present in 39 and 34 of the 52 MMR deficient patients in this cluster. s_AB_, the distance between the MMR deficient and MMR intact consensus networks in the interactome, was calculated as 0.1166. A positive s_AB_ value indicates the two networks do not overlap in the interactome and can be regarded as distinct molecular entities.

**Figure 1:**
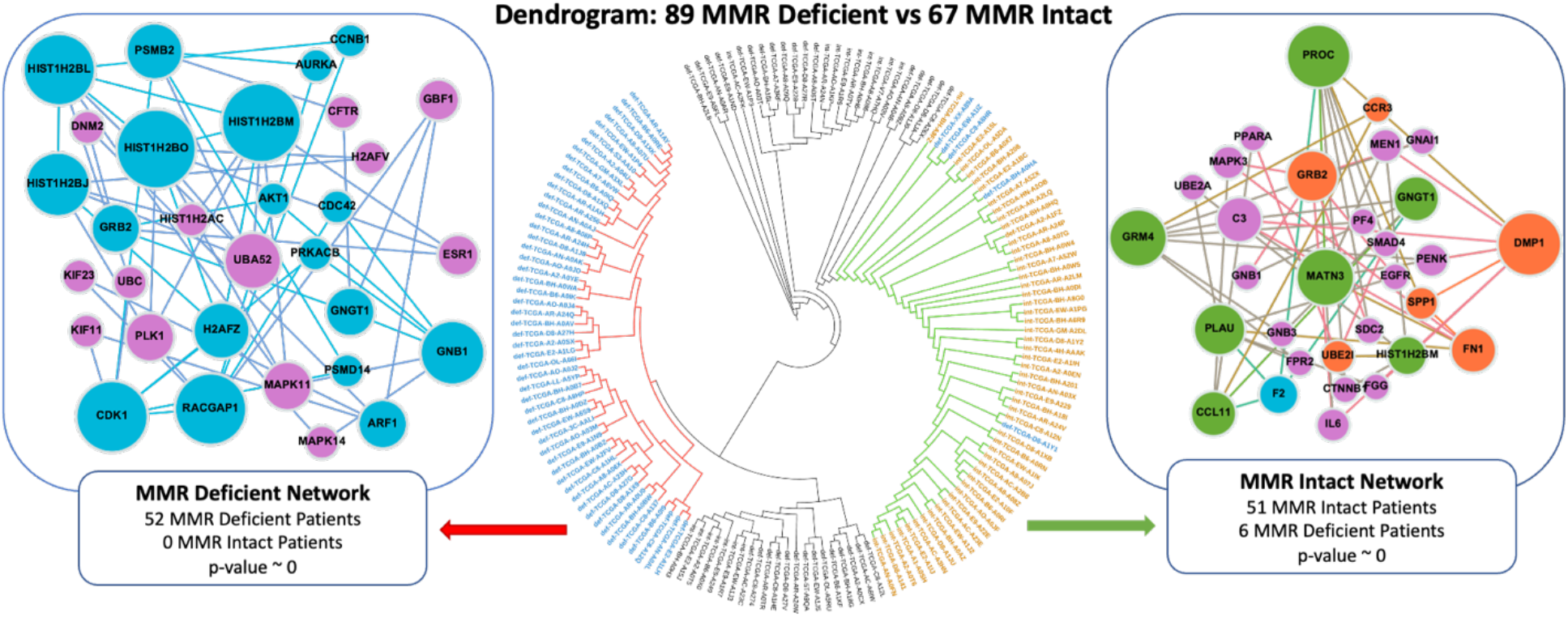
Dendrogram and consensus networks of MMR deficient (left) and MMR-intact specific clusters (right). Larger nodes indicate higher levels of connectivity. Patients with red branches are part of a significant cluster that create the MMR deficient consensus network (left). Patients with green branches contribute to the MMR intact consensus network (right). Orange nodes denote genes unique to MMR intact patients, blue nodes denote genes unique to MMR deficient patients, green nodes denote genes present in both MMR deficient and MMR intact networks, and purple nodes denote imputed genes. ITOL was used to generate the circular dendrogram^47^.

### Molecular and Clinicopathologic Profile

Molecular, clinical and pathologic findings were compared with MMR status on the cBioPortal. Overall, 907 patients were unaltered and 89 were altered. An overview of MMR mutations by protein is available in Supplemental Table 1 and overlapping MMR mutational signatures can be seen in Fig 2a. MMR deficiency was found to be associated with a higher mutation count (p<10e-10) and tumor mutational burden (p<10e-10), (Fig 2b); and a higher fraction genome alteration (p=5.077e-6), (Fig 2c). The results for OncoPrint events per patient can be seen in Fig 2d. Regarding breast cancer molecular subtypes, MMR deficiency was found to be associated with basal breast cancers (p=<10e-10) (Fig 2e). MMR deficiency status was found to be associated with increased rates of invasive ductal carcinoma compared to lobular and breast cancer of specific types (p=1.858e-3) (Fig 2f). Alterations in MMR were also found to be associated with higher MSI MANTIS scores (p=1.290e-5)(Fig 2g); as well as MSISensor scores (p=2.27e-10) (Fig 2h). All comparisons between MMR and the clinicopathological profile can be seen in Supplemental Table 2. There were no significant differences between progression-free, overall, disease-free, and disease-specific survival (p>0.05), (Fig 2 i-l). However, once patients were separated into individual MMR deficient groups, better overall survival was seen compared to MMR intact patients (P=0.00) (Fig 2m). HER2 enriched tumors were found to have worse disease-free survival (p=0.049), progression-free survival (p=0.723e-3), and disease-specific survival (p=0.029). While TN/basal subtype cancer had better overall survival (p=0.034). There was no clinical significance of MMR in Luminal A or B BC subtypes (Figure 3).

**Figure 2.**
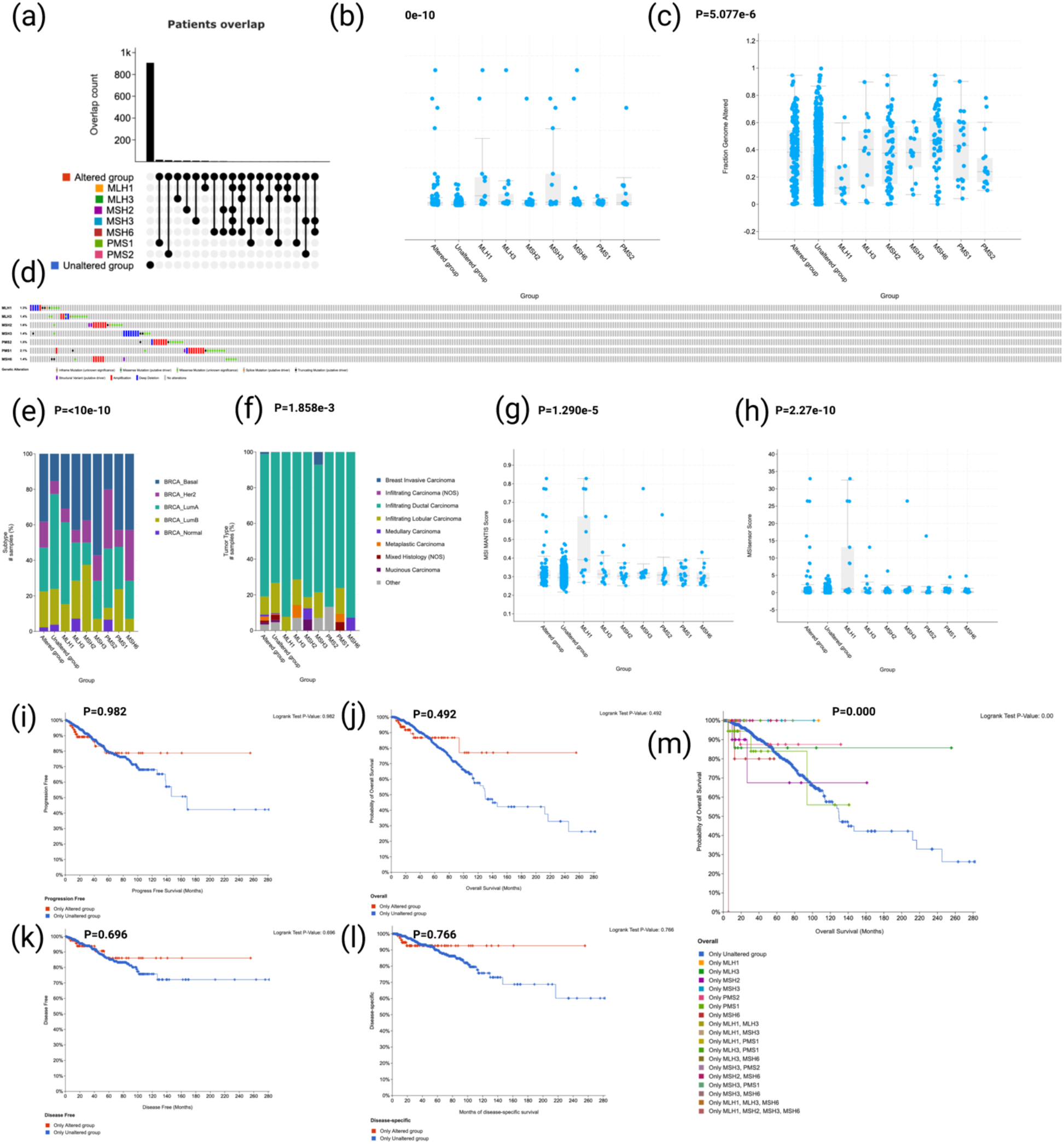
Molecular and clinicopathologic comparisons between MMR mutated and Intact groups. (a) MMR alteration groups with overlapping profiles; (b) MMR deficiency according to tumor mutational burden; (c) MMR deficiency according to fraction genome alteration; (d) OncoPrint events per patient; (e) MMR deficiency according to breast cancer molecular subtypes; (f) MMR deficiency according to breast cancer histologic subtypes; (g) Alterations in MMR in association with MSI MANTIS score; as well as MSISensor scores (h); (i) progression-free survival; (j) overall survival, disease-free survival (k), (l) disease-specific survival; and (m) overall survival according to individual MMR deficient groups.

**Figure 3.**
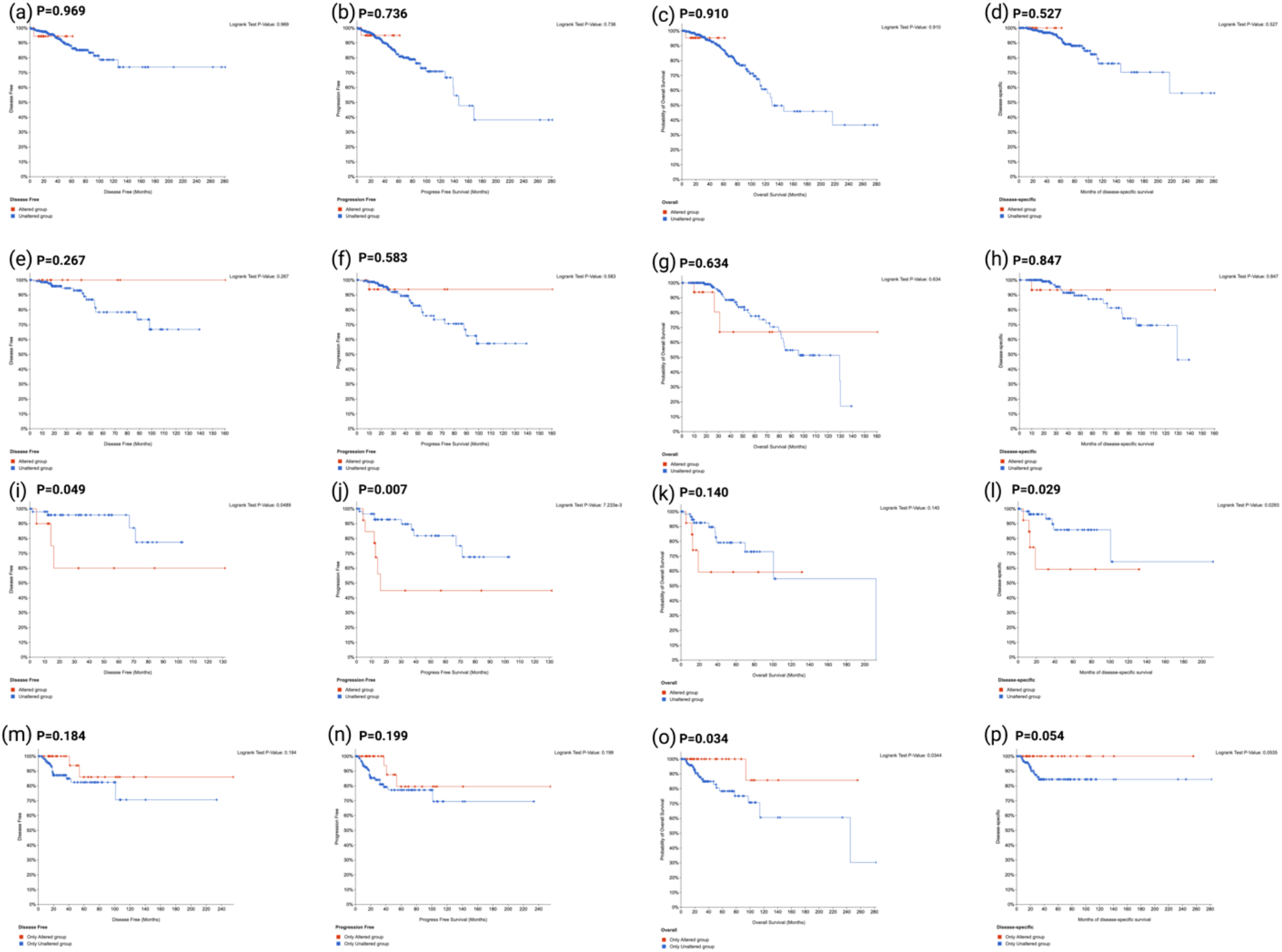
Survival analyses across different molecular breast cancer subtypes for MMR mutated tumors. (a) Luminal A: disease-free survival; (b) Luminal A: progression-free survival; (c) Luminal A: overall survival; (d) Luminal A: disease-specific survival; (e) Luminal B: disease-free survival; (f) Luminal B: progression-free survival; (g) Luminal B: overall survival; (h) Luminal B: disease-specific survival; (i) HER2: disease-free survival; (j) HER2: progression-free survival; (k) HER2: overall survival; (l) HER2: disease-specific survival; (m) Basal: disease-free survival; (n) Basal: progression-free survival; (o) Basal: overall survival; (l) Basal: diseasespecific survival.

### MMR Deficient Cohort C71

The C71 cohort included 52 of the 89 MMR deficient breast cancer patients. MSH2 had a somatic frequency of 7.7% with 1 driver mutation: 1 SV/fusion; and 3 VUS missense. MSH3 had a somatic mutation frequency of 7.7% with 1 driver mutation truncating, and 4 VUS, all of which were missense. MSH6 had a somatic mutation frequency of 11.5%: 2 driver mutations: 1 truncation and 1 SV/fusion; 6 VUS were seen, all missense. MLH1 had a somatic mutation frequency of 7.7% with 2 driver mutations: 1 missense and 1 spice; and 3 VUS, all of which were missense. PMS1 had a somatic mutation frequency of 13.5% with 2 driver mutations: 1 truncating and 1 SV/fusion; 5 VUS were seen, all missense. MLH3 had a somatic mutation frequency of 9.6% and contained 5 VUS, all of which were missense. PMS2 had a somatic mutation frequency of 9.6% and contained 2 driver mutation, all truncating: and 3 VUS, all missense. Overall, there were 10 driver mutations and 29 VUS, making VUS the majority (74%). Regarding molecular subtype, 28 were basal (TN), 9 HER2 enriched, 9 Luminal B and 6 luminal A. We found the C71 cohort to be more commonly associated with TN hormone receptor status (p=0.0004) as well as undergoing radiation therapy as part of their treatment course (p=0.0492).

The C71 cohort has good overall survival metrics with a 84.97% disease free survival, 75.37% progression free survival, 76.38% overall survival, and 90.71% disease specific survival (Supplemental Figure 2). Comparative analysis between the C71 MMR deficient cohort and control MMR deficient cohort can be seen in Supplemental Table 2.

## Discussion

Alterations in MMR mechanism has long been understood to cause cancer since knowledge gained from bacteria ultimately became relevant to human carcinogenesis in 1993 when the replication errors (RER) phenotype was discovered^25^. MMR is seen to be particularly relevant today in colorectal^26^ and endometrial^27^ carcinomas. Most recently, the use of immune checkpoint inhibitors has been approved and is seen to be appropriate in all MMR-deficient cancers^28^. However, in the clinical management of breast cancer, immune checkpoint inhibitors are used almost exclusively in patients with TN/basal biomarker status^29^.

The clinical findings in the present study support that MMR mechanisms are distinct depending on a tumor’s hormone receptor status and corresponding breast cancer molecular subtype. Fusco et al. demonstrated that breast cancer patients which were Luminal B-like and MMR deficient showed shorter overall survival than those who were MMR intact ^30^. On the other hand, they found patients with ER negative breast cancers treated with chemotherapy to live longer with MMR deficiency. We also found that TN/basal molecular status and MMR deficiency to be associated with improved survival. Some possible explanations for improved survival with MMR deficiency in TNBC is that (1) MMR deficiency was most common in these tumors, (2) immune check point inhibitors are commonly used in this cancer subtype, and (3) MMR deficient tumors have been shown to respond to checkpoint point inhibition across many cancer subtypes, although breast was not studied^31^. Cheng et al. found MMR deficiency to be significantly associated with worse overall and disease-specific survival in ER positive breast cancer patients who were all treated with tamoxifen as an isolated adjuvant systemic therapy^32^. However, there were only 31 MMR-deficient patients identified by IHC in the entire study^32^. In the present study, there were 89 MMR deficient patients and shorter survival was demonstrated in patients with HER2 molecular breast cancer status who were MMR deficient.

We showed histone related proteins as hub proteins in MMR deficient patients, particularly in cohort C71. DNA MMR is understood to be less efficient in targeting and replacing mispairs packaged in chromatin, as MMR must either compete for access to naked DNA before histone deposition or actively move nucleosomes^33^. Previous studies have demonstrated that neither purified MMR proteins nor nuclear extracts of human cells can repair DNA mismatches in the context of chromatin *in vitro*^34, 35^. Li et al. have further postulated that this may due to the chromatin histone structure itself inhibiting communication between the mismatch and nick site or that MutS may not efficiently recognize DNA mispairs when bound by a histone octamer^36^. Recently, it has been suggested that the MutSα interaction with the chromatin H3K36me3 active histone mark may also play a role in promoting MMR activity at sites of transcription^37^. Regarding the interactive complexity of the chromatin landscape, HUB2 gene signatures have been found to antagonize H3K27me3 marks in plant growth and developement^38^. HUB2 has been shown to increase histone H2B monoubiquitination in eukaryotes^39^. The MMR deficient C71 cohort appears to be a favorable predominantly ductal BC subtype associated with TN molecular status. The association with radiation status was likely secondary to the fact that these tumors were predominantly TN (54%), and TN status is associated with higher radiation therapy (RT) utilization at 69.2% compared to 55.7% in ER+/HER2+, 57.1% in ER+/HER2-, and 65.6% in ER-/HER2+ patients^40^. There was no difference between markers of microsatellite instability and other metrics of tumor mutational burden between MMR groups, or any difference in associations between overall and progression free survival.

Further studies examining MMR in relation to histone proteins and epigenetic modifications generally in TNBC could be beneficial. Recently histone lysine methyltransferases (KMTs) have emerged as attractive drug target in BC, although therapies targeting histone modifications (HMs) are still in the initial phases^41^. KMT nuclear receptor binding SET domain protein 2 (NSD2) has been shown to be overexpressed in TNBC tumors and control the expression of EGFR and ADAM9, a member of the ADAM (a disintegrin and metalloproteinase) family which releases growth factors including HB-EGF^42^. NSD2 may be identified as a major epigenetic regulator in TNBC.

The possible concordance between MMR by immunohistochemistry (IHC) and NGS may explain some discrepancies between the two previous studies and this study^30, 32^. Importantly, Fusco et al. showed a rate of discrepancy between IHC and molecular MSI analysis to be very high (91%)^30^. In the study the data was obtained, no definitive IHC was performed to examine protein expression. Validating IHC and its relationship to NGS results will be important for guiding diagnostic MMR workflows in BC. Current trends are moving away from traditional IHC and towards more comprehensive molecular profiling. Evaluation of any mutation detected by NGS in any of the 7 MMR deficiency genes found in our study may be appropriate. Patients with VUS should also be labelled as MMR deficient. The following is supported by distinct proteomic networks and clinically significant differences in survival for patients in the C71 cohort compared to the remaining MMR deficient patients. VUS defined most mutations (74%) in the C71 cohort.

In November 2020, the U.S. Food and Drug Administration (FDA) approved Keytruda (pembrolizumab) in combination with chemotherapy for unresectable locally advanced or metastatic triple-negative, PD-L1-positive breast cancers based on the results of the KEYNOTE-355 clinical trial^43^. More recently, MMR has been postulated as a molecular target in TNBC precision oncology harboring an increased sensitivity to immunotherapy^44^. MMR testing could be used in clinical practice to help guide treatment based on immunotherapy, particularly in HER2 enriched and TN tumors as we found these to be more proportionally involved by MMR. Also, mutational signatures associated with MMR have been found to be enriched in breast cancer brain metastases compared to primary breast tumors^45^. Suggesting there may be a role of MMR and PD-L1 testing in metastatic breast cancer specimens.

It is important to mention the numerous pitfalls in the present study including that we were unable to determine immune cell density and phenotypes in the tumoral microenvironment. MMR-deficient tumors often exhibit a high mutational burden and express neoantigens generated by frameshift mutations in coding microsatellites which stimulate lymphocytic infiltration as well as up-regulation of inflammatory cytokines^46^.

Finally, it would have been beneficial to validate the findings of our study in a second independent breast cancer cohort. Since the primary aim of our study is secondary data analysis but as a future reference it would also be beneficial to perform experimental validation of these findings.

In summary, the present study demonstrates MMR deficient BC to be a distinct molecular subtype with unique PPI networks involving histones hub genes and variable clinical significance depending on a patients individual HR status. Molecular subtyping based on MMR could be important for characterizing tumors as MMR deficient and guiding the use of immune checkpoint inhibition and other targeted therapies in both TN as well as HER2 enriched tumors.

## Data Availability

All data collected and analyzed in this study is available at cBioPortal (https://bit.ly/3g4lTM4).

## Author Contributions

SH, CC, YB, YW, AU, EDGU interpreted data, performed study concept and design, acquired the study materials, performed analysis, read, revised, and approved the final paper.

