## Supplementary material for "MMR Deficiency Defines Distinct Molecular Subtype of Breast Cancer with Unique Proteomic Networks and Variable Clinical Significance": Supplemental_Data.pdf

**Supplemental Figures**

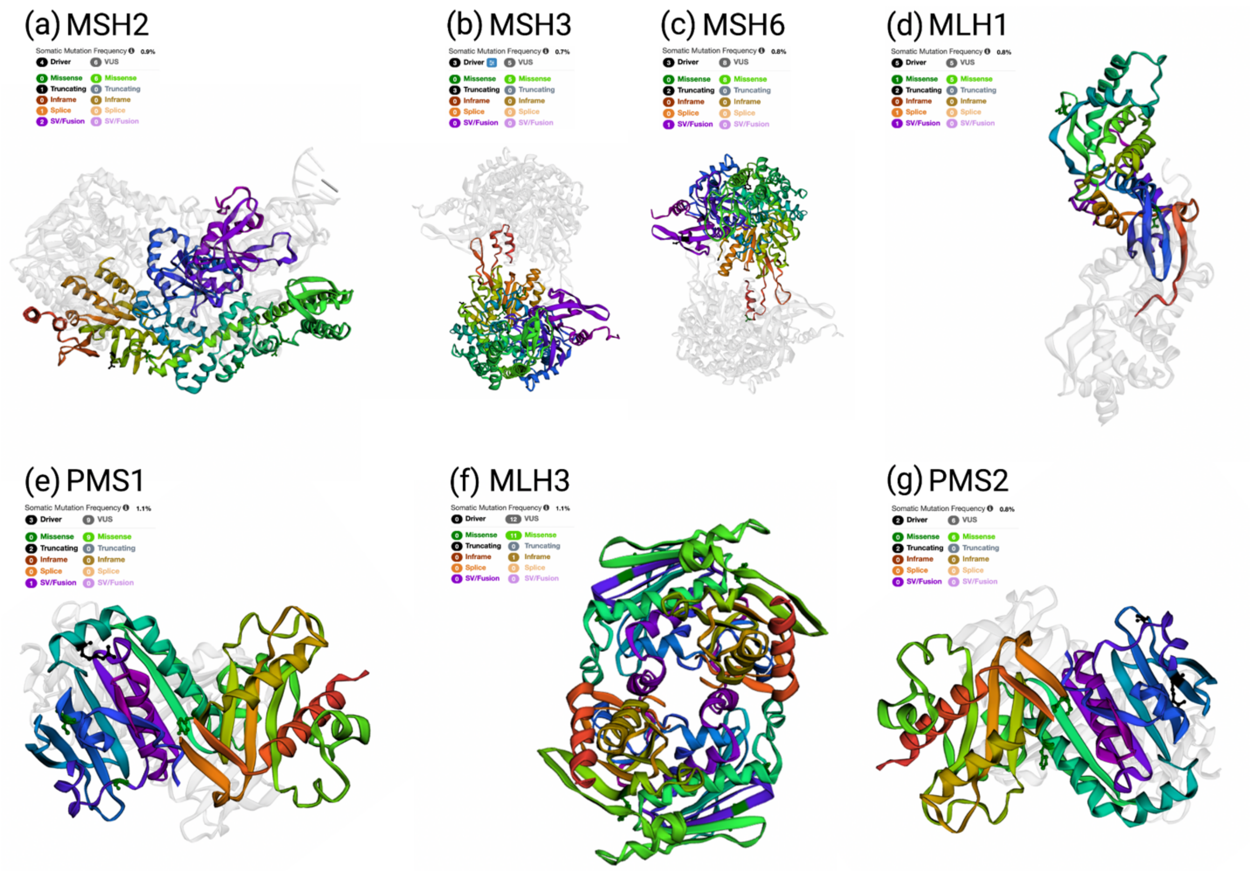


**Supplemental Figure 1. Landscape of MMR genomic mutations with corresponding 3D protein structural mutational findings.** (a) MSH2: PDB1ewq:crystal structure taq muts complexed with a heteroduplex DNA at 2.2a resolution, ChainA: dna mismatch repair protein muts (b) MSH3: PDB5x9w:mismatch repair protein, ChainB: dna mismatch repair protein muts; (c) MSH6: PDB5yk4:mismatch repair protein, ChainB:dna mismatch repair protein muts; (d) MLH1: PDB3rbn: crystal structure of mutl protein homolog 1 isoform ChainA: dna mismatch repair protein mlh; (e) PMS1: PDB1h7s: n-terminal 40kda fragment of human pms2, ChainA: pms1 protein homolog 2; (f) MLH3: PDB1b62:mutl complexed with adp ChainA: protein (mutl); (g) PMS2: PDB1h7s:n-terminal 40kda fragment of human pms2, ChainA: pms1 protein homolog 2.

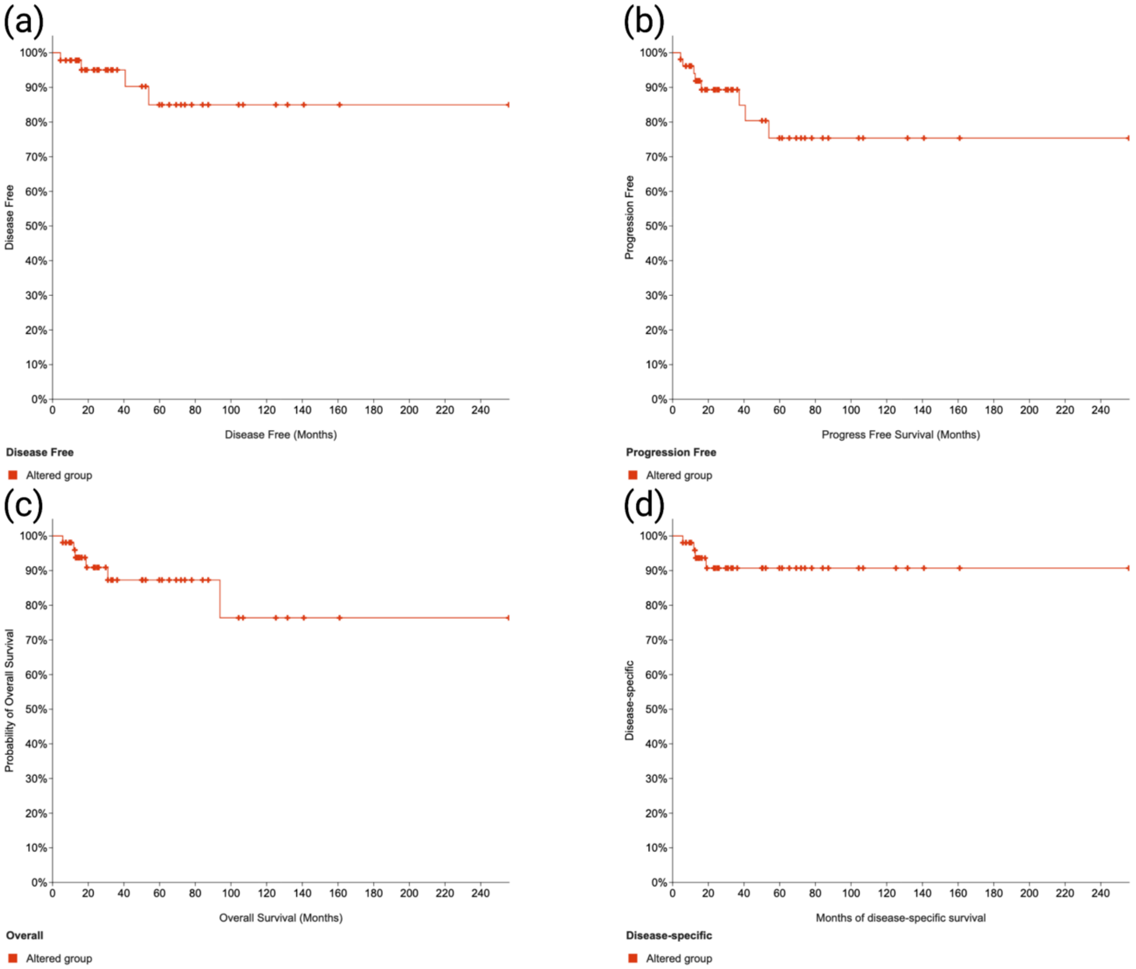


**Supplemental Figure 2. Survival analyses for the C71 MMR deficient cohort.** (a) 84.97% disease free survival, (b) 75.37% progression free survival, (c) 76.38% overall survival, and a (d) 90.71% disease specific survival.

### Supplemental Tables

**Supplemental Table 1. Overview of MMR mutations by protein.**

| MMR Protein | Patients (89 Total) |
|---|---|
| PMS1 | 18 |
| PMS2 | 14 |
| MLH3 | 10 |
| MSH2 | 10 |
| MSH3 | 9 |
| MLH1 | 8 |
| MSH6 | 5 |
| MSH2, MSH6 | 5 |
| MLH1, MSH2, MSH3, MSH6 | 1 |
| MLH1, MLH3, MSH6 | 1 |
| MSH3, PMS1 | 1 |
| MLH1, MSH3 | 1 |
| MLH3, MSH6 | 1 |
| MLH1, PMS1 | 1 |
| MLH1, MLH3 | 1 |
| MLH3, PMS1 | 1 |
| MSH3, PMS2 | 1 |
| MSH3, MSH6 | 1 |

**Supplemental Table 2.** Comparisons between MMR deficient and intact patients.

| Variable | Statistical Test | P-Value | Q-Value |
|---|---|---|---|
| Mutation Count | Wilcoxon Test | **$< 10^{-10}$** | $< 10^{-10}$ |
| TMB (nonsynonymous) | Wilcoxon Test | **$< 10^{-10}$** | $< 10^{-10}$ |
| Buffa Hypoxia Score | Wilcoxon Test | **2.16e-9** | 3.45e-8 |
| Subtype | Chi-squared Test | **8.65e-9** | 1.04e-7 |
| Winter Hypoxia Score | Wilcoxon Test | **2.55e-8** | 2.45e-7 |
| Ragnum Hypoxia Score | Wilcoxon Test | **4.00e-7** | 3.041e-6 |
| MSIsensor Score | Wilcoxon Test | **4.43e-7** | 3.041e-6 |
| Fraction Genome Altered | Wilcoxon Test | **4.765e-4** | 2.859e-3 |
| MSI MANTIS Score | Wilcoxon Test | **1.393e-3** | 7.430e-3 |
| Oncotree Code | Chi-squared Test | **2.374e-3** | 0.0104 |
| Cancer Type Detailed | Chi-squared Test | **2.374e-3** | 0.0104 |
| Primary Lymph Node Presentation Assessment | Chi-squared Test | **3.731e-3** | 0.0149 |
| Tumor Type | Chi-squared Test | **0.0144** | 0.0532 |
| International Classification of Diseases for Oncology, Third Edition ICD-O-3 Site Code | Chi-squared Test | **0.0499** | 0.171 |
| Radiation Therapy | Chi-squared Test | 0.0586 | 0.188 |
| ICD-10 Classification | Chi-squared Test | 0.0832 | 0.250 |
| Aneuploidy Score | Wilcoxon Test | 0.133 | 0.336 |
| Form completion date | Chi-squared Test | 0.283 | 0.603 |
| American Joint Committee on Cancer Tumor Stage Code | Chi-squared Test | 0.285 | 0.603 |
| Last Communication Contact from Initial Pathologic Diagnosis Date | Wilcoxon Test | 0.289 | 0.603 |
| Tissue Source Site Code | Chi-squared Test | 0.319 | 0.612 |
| Tissue Source Site | Chi-squared Test | 0.319 | 0.612 |
| Race Category | Chi-squared Test | 0.385 | 0.708 |
| New Neoplasm Event Post Initial Therapy Indicator | Chi-squared Test | 0.439 | 0.708 |
| In PanCan Pathway Analysis | Chi-squared Test | 0.443 | 0.708 |
| Tissue Prospective Collection Indicator | Chi-squared Test | 0.457 | 0.708 |
| Tissue Retrospective Collection Indicator | Chi-squared Test | 0.457 | 0.708 |
| Other Patient ID | Chi-squared Test | 0.485 | 0.728 |
| Person Neoplasm Cancer Status | Chi-squared Test | 0.556 | 0.808 |
| Sex | Chi-squared Test | 0.608 | 0.842 |
| Neoplasm Disease Lymph Node Stage American Joint Committee on Cancer Code | Chi-squared Test | 0.614 | 0.842 |
| American Joint Committee on Cancer Metastasis Stage Code | Chi-squared Test | 0.725 | 0.966 |
| Ethnicity Category | Chi-squared Test | 0.779 | 0.985 |
| Birth from Initial Pathologic Diagnosis Date | Wilcoxon Test | 0.828 | 0.992 |
| International Classification of Diseases for Oncology, Third Edition ICD-O-3 Histology Code | Chi-squared Test | 0.889 | 0.992 |
| Diagnosis Age | Wilcoxon Test | 0.905 | 0.992 |

| | | | |
|---|---|---|---|
| American Joint Committee on Cancer Publication Version Type | Chi-squared Test | 0.908 | 0.992 |
| Neoplasm Disease Stage American Joint Committee on Cancer Code | Chi-squared Test | 0.930 | 0.992 |
| Neoadjuvant Therapy Type Administered Prior To Resection Text | Chi-squared Test | 0.958 | 0.993 |
| Prior Diagnosis | Chi-squared Test | 0.989 | 0.993 |

**Supplemental Table 3.** Comparisons between MMR deficient cohort C71 and control cohort.

| Patient Characteristics | C71 Cohort (n =52) | Control Cohort (n = 37) | p value |
|---|---|---|---|
| Age | | | *0.125 |
| Mean | 56.5 | 61.1 | |
| Stage (AJCC) | | | **0.353 |
| 1 | 9 | 3 | |
| 2 | 30 | 26 | |
| 3-4 | 12 | 7 | |
| Histologic type | | | **0.068 |
| Ductal | 48 | 29 | |
| Not ductal | 4 | 8 | |
| Radiation Therapy | | | ****0.0492** |
| Yes | 28 | 11 | |
| No | 24 | 24 | |
| Molecular subtype | | | ***+0.0004** |
| Luminal A | 6 | 16 | |
| Luminal B | 9 | 9 | |
| HER2 | 9 | 4 | |
| Basal (TN) | 28 | 6 | |
| Aneuploidy Score | | | *0.514 |
| Mean | 13.80 | 12.75 | |
| Fraction Genome Altered | | | |
| Mean | | | |
| TMB | | | *0.708 |
| Mean | 11.57 | 13.98 | |
| MSI MANTIS Score | | | *0.413 |
| Mean | 0.33 | 0.34 | |
| MSI sensor Score | | | *0.661 |
| Mean | 1.89 | 2.42 | |
| Mutation Count | | | *0.687 |
| Mean | 341.21 | 418.73 | |
| Overall Survival Status | | | **0.730 |
| Yes | 46 | 34 | |
| No | 6 | 3 | |
| Progression Free Status | | | **0.349 |
| Yes | 44 | 34 | |
| No | 8 | 3 | |

*Unpaired t test; **Fisher's exact test; *+$\chi 2$ contingency test.
